# Adapting flow cytometry for studying immune cell seasonality in wild migratory bats

**DOI:** 10.64898/2026.02.03.702504

**Authors:** Meagan Allira, Kristin E. Dyer, Bret M. Demory, Mackenzie G. Hightower, Beckett L. Olbrys, Kaylee M. Norman, Mark L. Lang, Daniel J. Becker

**Affiliations:** School of Biological Sciences, University of Oklahoma, Norman, OK, USA; Department of Microbiology and Immunology, University of Oklahoma Health Campus, Oklahoma City, OK, USA; Department of Microbiology and Immunology, Roy J. and Lucille A. Carver College of Medicine, University of Iowa, Iowa City, Iowa, USA

**Keywords:** cellular immunity, ecoimmunology, Mexican free-tailed bats, *Tadarida brasiliensis*, One Health

## Abstract

Identifying the drivers of wildlife immunity is critical for assessing stressor impacts and zoonotic risks. However, such studies are limited by logistical challenges of wildlife research and lack of species-specific reagents. We adapt flow cytometry, typically confined to laboratory settings, to field settings to profile cellular immunity with small blood volumes and extended sample holding times. We apply these methods to analyze immune cell seasonality in migratory Mexican free-tailed bats (*Tadarida brasiliensis*). We confirmed four antibodies recognizing CD3, CD79a, MHCII, and CD11b that were originally validated in Egyptian fruit bats (*Rousettus aegyptiacus*), allowing us to quantify T and B cells, macrophages, and neutrophils, respectively. Flow cytometry outperformed hematology in quantifying leukocyte profiles and revealed pronounced immune cell seasonality. Adaptive cells steadily increased between spring and fall migration. Neutrophils were most abundant during the reproductive period and decreased during migrations, whereas B cells were most abundant after reproduction and before fall migration; granulocytes as a whole, macrophages, and T cells had no seasonality. Females had more B cells than males but did not differ in other cells. Our findings lay the groundwork for applying flow cytometry to field studies of wildlife and provide important insights into the seasonality of bat immunity.

## Introduction

Wildlife rely on diverse immune strategies to manage infections (Pedersen and Babayan 2011), which can be altered through human impacts on natural systems (Martin et al. 2010). Such disturbances can destabilize disease dynamics by degrading landscape immunity (i.e., the suite of ecological conditions within an ecosystem that collectively maintain and strengthen wildlife immune function), ultimately increasing opportunities for spillover from wildlife to humans (Plowright et al. 2021). Renewed attention to cross-species transmission following the COVID-19 pandemic has suggested that environmental changes can not only modify contact between humans and wildlife, but also alter the immune state of wild hosts and increase their susceptibility to infection and duration or intensity of pathogen shedding (Becker et al. 2020a; Plowright et al. 2024). As human expansion continues, better characterizing and identifying the drivers of wildlife immune fluctuations will be increasingly important, particularly for species host to zoonotic pathogens. Understanding seasonality in immunity will further our ability to predict when different species are likely to shed pathogens, informing disease risk management.

Across taxa, animals mitigate energetically demanding life stages through trade-offs among costly physiological functions (Zera and Harshman 2001; Martin et al. 2008). Given finite energetic resources, hosts often rely on allocating resources in one trait that requires a decrease in another, such as between reproduction and longevity (Cox et al. 2010). This principle also extends to the immune system, both between these functions and immunity (Ardia 2005) and between different branches of the immune system (Minias et al. 2023). The direction of such trade-offs is largely governed by relative costs of innate and adaptive immunity. Although these two systems are interconnected, with innate responses activating adaptive pathways to resolve infection and minimize tissue damage (Clark and Kupper 2005; Iwasaki and Medzhitov 2015), they also specialize in distinct modes of defense. Innate immunity, led by macrophages and neutrophils, forms broad and rapid defense by recognizing pathogen-associated molecular patterns and triggering cytokine cascades (Medzhitov and Janeway 2000; Akira et al. 2006). By contrast, adaptive immunity targets pathogens through antigen-specific responses mediated by cytotoxic T cells, which target and eliminate infected cells, and helper T cells, which stimulate B cell production to generate antibodies and generate memory for long-term immunity (Harty et al. 2000; Zhu et al. 2010; Cyster and Allen 2019). As such, innate immunity has low developmental costs, but it is energetically costly to both activate and maintain, with unchecked responses also causing systemic inflammation and damaging host tissues. By contrast, adaptive immunity requires higher developmental investment but provides efficient, low-cost responses upon secondary exposure (Rauw 2012; McDade et al. 2016). These relative costs mean that trade-offs can manifest as hosts relying on adaptive immunity while resources are plentiful but instead prioritizing innate immunity during energetically demanding periods to conserve resources.

Intrinsic and extrinsic stressors, including but not limited to periods of reproductive activity, food limitation, and temperature extremes, can all force trade-offs between and possibly within innate and adaptive immunity (Norris and Evans 2000; Lochmiller and Deerenberg 2000; Martin et al. 2008). Long-distance migration, defined as the seasonal movement between breeding and wintering grounds, represents a particularly high-cost intrinsic stressor (Wikelski et al. 2003; Dingle 2014). Migration can modulate immunity in ways that alter host susceptibility to new or chronic infections, and reshape disease dynamics (Altizer et al. 2011; Becker et al. 2020b). Most studies of migration and immunity have focused on birds, where innate and sometimes adaptive immunity can be suppressed during migration (Owen and Moore 2008; Nebel et al. 2012; Eikenaar and Hegemann 2016). However, despite this behavior being common in several bat families (Fleming et al. 2003; Bisson et al. 2009), the effects of migration on immunity in bat hosts remains poorly understood (Gonzalez et al. 2024). This is a notable knowledge gap, given that many bat species can tolerate otherwise virulent infections (Cummings et al. 2025), in part through unique immune mechanisms such as dampened inflammatory responses and other mechanisms (Ahn et al. 2019; Irving et al. 2021; Morales et al. 2025). On the one hand, this tolerance of infection could allow bats to readily disperse pathogens over large spatial scales during migration (Peel et al. 2013). Yet on the other hand, immunological trade-offs experienced during migration, thus far observed in a small number of bat case studies (Voigt et al. 2020; Rogers et al. 2022), could modify bat defense in ways that facilitate susceptibility to infection or greater shedding of otherwise chronic infections. Better characterizing the role of these seasonal movements on bat immune function is key to understanding the broader consequences of migration on zoonotic risk (Altizer et al. 2011).

However, our ability to characterize the immune systems of wild bats is limited by a lack of species-specific reagents, field-friendly assays, and more targeted tools to examine specific immune cells (Baker et al. 2013; Schountz 2014; Banerjee et al. 2020; Irving et al. 2021).Studying cellular immunity in wildlife, and in bats especially, has been mostly limited to using hematology to measure total and differential white blood cell counts (Schneeberger et al. 2013; Voigt et al. 2020), which broadly distinguishes lymphocytes, monocytes, and granulocytes (i.e., neutrophils, eosinophils, and basophils) but provides limited resolution. Flow cytometry instead enables finer characterization of immune cells in tissues such as blood, including B and T cell subsets (Adan et al. 2017; McKinnon 2018), but has been largely confined to laboratory or captive systems (Sylvester et al. 2018; Hunka et al. 2020) due to needs for large blood volumes and species-specific reagents for staining cells. However, studies have begun adapting antibody panels for bats, although this work has primarily focused on model captive breeding colonies (Wang et al. 2021), mostly Old World fruit bats (i.e., Pteropodidae) (Martínez Gómez et al.2016; Periasamy et al. 2019; Gamage et al. 2020; Friedrichs et al. 2022; Chen et al. 2024). While this has set the stage for expansion of flow cytometry into field studies of bat cellular immunity, barriers remain to implement flow cytometry in wild systems. This includes but is not limited to testing antibodies in other bat taxa and use of smaller blood volumes, which is especially relevant for bat fieldwork due to blood collection limitations while avoiding lethal sampling.

Here, we adapt flow cytometry protocols for use in fieldwork settings to enable more precise characterization of wildlife cellular immunity. We first test protocols in laboratory mice to identify the smallest blood volume feasible to collect while still producing adequate fluorescence through flow cytometry. We then test antibodies previously validated in the Pteropodidae in wild, migratory Mexican free-tailed bats (*Tadarida brasiliensis*, family Molossidae). Using this antibody panel, we then compared immune cell counts from flow cytometry with those derived from blood smears to assess the accuracy of manual counts. Lastly, we test hypotheses about seasonality and trade-offs in cellular immunity in our migratory bat system by applying our antibody panel to blood samples collected across the occupancy period in western Oklahoma, capturing energetic stressors of spring migration, pregnancy, and preparation for fall migration. *Tadarida brasiliensis* is an ideal system for studying seasonality in immunity and such trade-offs due to its long migratory movements, large colony sizes, and exposure to diverse pathogens (Glass 1982; Ganow et al. 2015; Becker et al. 2024; Becker et al. 2025b).Given the energetic costs of both migration and reproduction, we predicted that bats would favor innate immune investment during these periods and shift allocation to adaptive responses during non-reproductive and non-migratory periods, thus reflecting life-history trade-offs in immunity.

## Methods

### Biology of the migratory bat system

Mexican free-tailed bats are an abundant, widespread, and partially migratory North American bat species that serve important roles in predating insects, with an estimated contribution of $12.2 million in pest control annually in the southwestern United States (Wiederholt et al. 2017). Many bats overwinter in Mexico in small, dispersed populations, with mating occurring prior to spring migration into the southwestern United States. One major migratory pathway includes Texas, Oklahoma, and Kansas, where females and males aggregate in maternity and bachelor colonies that can reach hundreds of thousands to tens of millions of bats during summer months (Bernardo and Cockrum 1962; Glass 1982; Wiederholt et al. 2013). Oklahoma hosts a small number of maternity and bachelor colonies in the western part of the state, with these cave roosts being fully occupied by May (Caire et al. 1989; Ganow et al. 2015). Here, females advance their pregnancy until giving birth in June and July, followed by pup rearing and eventual migration back to Mexico in September and October (Figure 1). We focused our analyses on the Selman Bat Cave and Alabaster Caverns State Park, two cave roosts separated by 8 km and representing a maternity and bachelor colony, respectively. The Selman Bat Cave holds up to approximately 50,000 Mexican free-tailed bats during summer (Ganow et al. 2015) and, in contrast to some populations in Texas (Stepanian and Wainwright 2018), includes only migratory bats. Alabaster Caverns State Park hosts a smaller Mexican free-tailed bat population alongside multiple other bat species (Caire et al. 1984; Roistacher et al. 2025). Our ongoing movement monitoring of Mexican free-tailed bats shows high connectivity of these caves (Dyer et al., unpublished), such that we included sampling at both sites to provide an even sex ratio. Prior banding and molecular analyses of these populations have shown the bats can migrate as far south as Veracruz, Mexico, a distance of 1,800 km (Glass 1982; Russell et al. 2005; Nichols et al. 2019). We can therefore infer that most bats arrive in the study region in spring following a long-distance migration.

**Figure 1.**
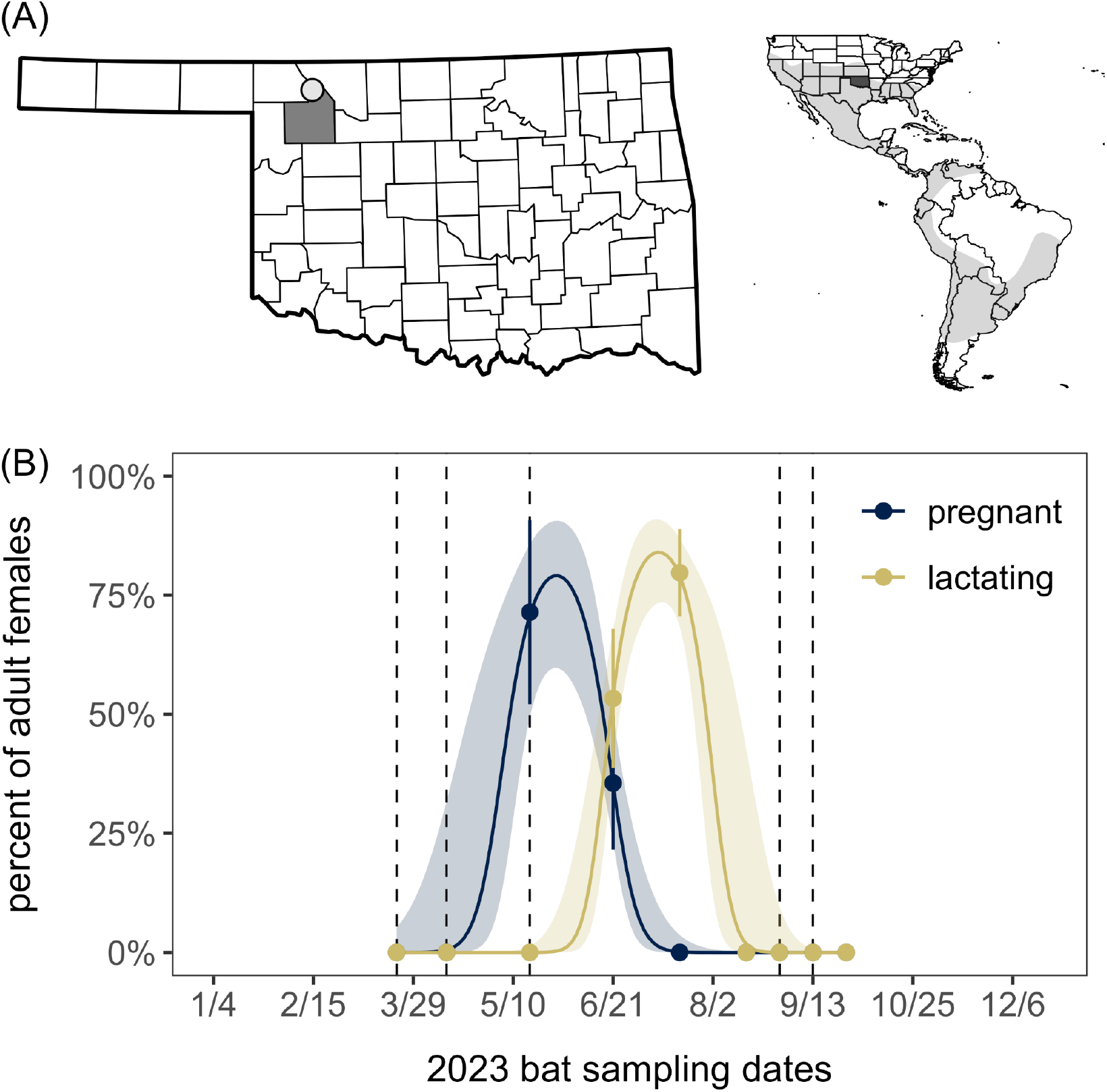
(A) Study site in Woodward County, Oklahoma, relative to the Mexican free-tailed bat geographic range, as defined by the International Union for Conservation of Nature. (B) Sampling timepoints for flow cytometry (dashed lines) relative to the bat migratory and reproductive cycle, based on a larger sample of adult females captured in 2023 (*n* = 314).Seasonal proportions of pregnant and lactating adult females are shown with 95% confidence intervals (Wald intervals). Overlaid are fitted values and 95% confidence intervals from generalized additive models of each reproductive state (binary response) with a cyclic cubic smooth term of week (knots were bound within weeks 1 and 52), fit using the *mgcv* package in R and restricted maximum likelihood (setting γ=1.4 to reduce overfitting risks) (Wood 2017).

### Optimizing blood volumes

Traditional flow cytometry uses a minimum of 50 uL blood from model organisms such as mice (Dertinger et al. 2004). However, laboratory mice weigh more (approximately 25 grams) than most North American bat species, including Mexican free-tailed bats (on average, 11 grams).While 50 uL blood is a safe volume (i.e., less than 1% body mass) to collect from laboratory mice and similarly sized or larger bat species, this volume can be prohibitive for the large number of smaller bat species (Sikes and Gannon 2011; Becker et al. 2025c). Further, while such blood volumes are logistically tractable in laboratory settings, where animals are well-fed and hydrated, 50 uL blood can be challenging to collect in field conditions. As such, we performed an initial optimization of protocols using laboratory mice to reduce blood volumes needed.

We determined the smallest blood volume required to produce quality fluorescence while also maintaining accurate cell ratios and thus provide precise quantification of target cell populations. We collected 100 µL blood from the retro-orbital vein of mice (*n* = 2) into heparinized capillary tubes. Blood from both mice was pooled into tubes with 50 µL EDTA and then aliquoted into 50 µL, 40 µL, 30 µL, 20 µL, and 10 µL volumes. We then centrifuged samples (2000 rpm for five minutes) to remove EDTA before resuspending pellets in 200 µL 1X PBS. We next added 1:200 2.4G2 (Fc-region block to inhibit non-specific antibody binding; Biocell BE0307) and incubated samples at room temperature for five minutes. This monoclonal antibody has been used in previous *in vitro* studies of the black flying fox (*Pteropus alecto*) (Beauregard 2020), and Fc receptors of bats, mice, and humans have been shown to display relatively high sequence homology (Toshkova et al. 2023). We added 6 µL antibody cocktail (1.5 µL per color) to each sample for detection of APC-Cy7 (1:200 for CD19; Molecular Probes A15391), Brilliant Violet 450 (1:200 for CD4; BioLegend 103040), and PE (1:200 for Ly-6G; Tonbo Biosciences 50-5931). We used CD4 stained with 1:200 APC-Cy7, Brilliant Violet 450, and PE for single-color controls. Samples were then incubated (light protected) at room temperature for 45 minutes. We then washed cells three times with 1 mL of 1X PBS, incubating samples (light protected) for five minutes at room temperature between washes. After the final wash, 1 mL 1X FACS Lysing Solution was added for up to two washes per sample to remove red blood cell debris, incubating for five minutes between washes. Samples were then fixed in 1% paraformaldehyde for 20 minutes before analysis, after which they were then analyzed on a Stratedigm-3 (Stratedigm S1200Ex) flow cytometer at the University of Oklahoma (OU) Health Campus Flow Cytometry and Imaging Core to measure presence of fluorescently labeled cells.

### Antibody panel testing in Mexican free-tailed bats

We selected a subset of antibodies earlier validated in Egyptian fruit bats (*Rousettus aegyptiacus*) (Friedrichs et al. 2022) to test in Mexican free-tailed bats (Table 1). Initial antibody panel testing was performed with 30–40 µL blood from eight wild bats in March–April 2023 (see *Flow cytometry of wild bat blood leukocytes* below), as we determined this volume to be sufficient in our mouse trial (Figure S1). We tested the monoclonal antibody for CD3 in May 2023. All bat samples were prepared in the field, using the methods described above and developed during our mouse trials. Blood samples were stained with 1:200 PE (CD206; BD 566884), Brilliant Violet 605 (MHCII; BioLegend 101205), FitC (CD11b; BioLegend 101205), Pacific Blue (CD3; BioRad MCA1477PB), and APC (CD79a; BioLegend 333505). Samples were fixed and stored at 4°C, capped, and light protected during transport from the field to the OU Health Campus Flow Cytometry and Imaging Core. We found adequate fluorescence for CD3, CD11b, MCHII, and CD79a, while CD206 was incompatible with Mexican free-tailed bats (Table 1). Sample preparation and flow cytometry were performed by one individual (MA).

**Table 1.** Antibodies tested in Mexican free-tailed bats (*Tadarida brasiliensis*) for flow cytometry analysis, following their previous validation in the Egyptian fruit bat (*Rousettus aegyptiacus*).

| Antigen | Cell target | Clone | Species | Isotype | Fluorochrome | Compatible with<br>*Tadarida brasiliensis* |
| --- | --- | --- | --- | --- | --- | --- |
| CD3 | T cell | CD3-12 | Human | Rat IgG1 | Pacific Blue | Yes |
| CD11b | Neutrophil,<br>monocyte | M1/70 | Human,<br>mouse | Rat<br>IgG2b κ | APC | Yes |
| MHCII | Dendritic,<br>macrophage | 2G9 | Mouse | Rat<br>IgG2b κ | Brilliant Violet<br>605 | Yes |
| CD206 | Granulocyte<br>precursor | 15-2 | Human | Mouse<br>IgG1 κ | PE | No |
| CD79a | B cell | HM47 | Human | Mouse<br>IgG1 κ | FITC | Yes |

### Gating strategy for flow cytometry

Flow cytometry data acquisition was performed using CellQuest (BD), collecting a minimum of 50,000 events. We performed gating and analyzed dot plot cytograms using FlowJo. Forward scatter (FSC) and side scatter (SSC) parameters were used to determine cell size and internal complexity (Figure 2). Gates were established first around single cells, then around lymphocyte populations, which are smaller in size (i.e., lower on FSC) and have simple internal structure (i.e., lower on SSC), and granulocyte populations (higher on FSC and SSC). Positive and negative compensation beads (BioLegend 750002761 and 75002764) were used in accordance with manufacturer instructions for single-color controls, and compensation was established for each antibody (CD11b neutrophils, CD3 T cells, CD79a B cells, MHCII macrophages) by comparing peaks in sample fluorescence to those of the unstained cells and single-color controls.

**Figure 2.**
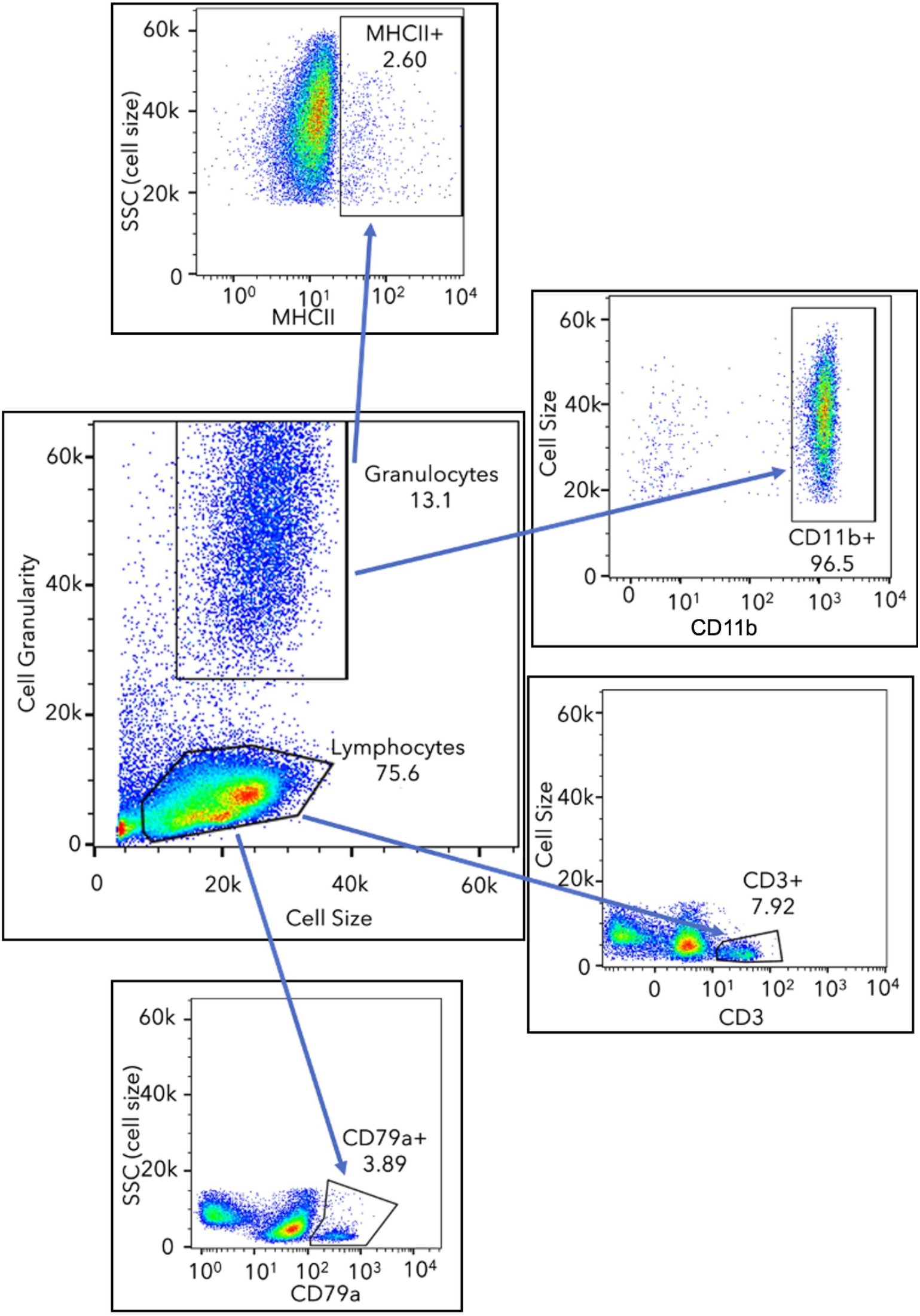
Representative cytograms, showing gating strategy of leukocyte populations based on size (FSC) and internal complexity (SSC) for MHCII+ (macrophages), CD11b+ (neutrophils), CD3+ (T cells), and CD79a+ (B cells).

### Flow cytometry of wild bat blood leukocytes

As part of long-term studies on immunity and infection in migratory Mexican free-tailed bats (Becker et al. 2024; Roistacher et al. 2025; Simonis et al. 2025), we captured bats using hand nets and mist nets at the Selman Bat Cave and Alabaster Caverns State Park at approximately monthly intervals between March and September 2023. We collected blood from the propatagial vein with sterile 27G needles and heparinized capillary tubes. Bats were held for an average of three hours prior to blood collection (± 0.1 hours). For flow cytometry, blood was deposited into 1.5 mL Eppendorf tubes and washed with 1X PBS. Samples were stained with the same antibodies used in our validation trials (see above) and fixed with 1% paraformaldehyde in the field, using the laboratory methods we adapted above with laboratory mice. However, to allow for intracellular staining of CD79a, from May onward (representing 73% of blood samples), Cyto-Fast Fix/Perm Buffer (BioLegend 426803) was instead used for fixation and permeabilization of cells and incubated for 20 minutes at room temperature. All centrifugation steps used a variable speed microcentrifuge (Gyro Plus, MiniPCR Bio). Samples were then held at 4°C for an average of 36 hours (± 4 hours) using a portable, nine-liter refrigerator (F40C4TMP) before being transported to the OU Health Campus Flow Cytometry and Imaging Core for analysis. To compare leukocyte profiles from flow cytometry with those from traditional hematology, we prepared blood smears on glass slides stained with Wright–Giemsa (Quick III, Astral Diagnostics). All bats were released at the capture site after processing.

### Hematological comparison with flow cytometry

We estimated leukocyte profiles from blood smears by identifying 100 leukocytes and recording the relative abundance of each type of white blood cell under 1000× magnification via oil immersion (Roistacher et al. 2025). Slides were read in duplicate by two users (KED and BLO). To compare these counts to those via flow cytometry, cell proportions were made comparable among methods. From hematology data, we defined granulocytes as the summed proportions of neutrophil, basophil, and eosinophil counts. Proportions were comparable between users for both granulocytes (ρ = 0.84, *p* < 0.001) and lymphocytes (ρ = 0.73, *p* < 0.001), such that we averaged proportions per bat between both reads. For flow cytometry, we calculated proportions of lymphocytes and granulocytes by dividing counts by total singlet cell counts. We then used Spearman correlations to compare granulocyte and lymphocyte proportions between methods.

### Analysis of seasonality and trade-offs in wild bats

We used our flow cytometry data to analyze seasonal changes and sex differences in immune cells alongside trade-offs among arms of the cellular immune system. To assess seasonality, we used the *mgcv* package in R to fit generalized additive models (GAMs), which can flexibly capture temporal trends (Wood 2017). We modeled each cell type as a quasibinomial response to account for overdispersion. For lymphocytes and granulocytes, we used these cell counts relative to total singlet cells, whereas we modeled B and T cells against total lymphocyte counts as well as macrophages and neutrophils against total granulocyte counts; this approach allowed us to model cellular proportions while accounting for variation in denominators (i.e., total singlet cells and total granulocytes or lymphocytes). We used a cyclic cubic spline of week to model seasonal trends, with knots bound within weeks 1 and 52. Each GAM also included the time between capture and sample collection (i.e., holding time) and between sample collection and flow cytometry analysis (i.e., processing time) as precision covariates (Laubach et al. 2021). Because one bat lacked data on holding time, we used the *missRanger* package to impute missing values using chained random forests (*n* = 1,000 trees) (Stekhoven and Bühlmann 2012; Wright and Ziegler 2017). All GAMs were fit using a logit link and restricted maximum likelihood, setting γ=1.4 to reduce risk of overfitting. Given our sample size, we also fit a separate set of analogous GAMs to assess sex differences in cellular immunity. To lastly evaluate possible trade-offs between these cell proportions, we used Spearman correlations on our individual-level data.

## Results

To limit unnecessary sampling of bats, we first used laboratory mice and their specific antibody panels to develop approaches for subsequent optimization in the field. We found fluorescence was adequate in blood volumes above or equal to 30 µL (Figure S1), which allows sampling small mammals while maintaining blood volume to less than 1% body weight. We then tested a subset of antibodies validated for *Rousettus aegyptiacus* in Mexican free-tailed bats, confirming four (CD3, CD79a, CD11b, and MHCII) of five monoclonal antibodies (Table 1). These initial analyses thereby allowed us to compare hematology to flow cytometry and to analyze seasonal fluctuations in immune cell populations within and between innate and adaptive immunity.

Between March and September 2023, we collected 30–40 µL blood from 37 Mexican free-tailed bats for flow cytometry, with paired blood smears from 30 individuals to compare methodology (Figure 3). Lymphocyte proportions were moderately but significantly correlated between hematology and flow cytometry (ρ = 0.62, *p* < 0.001). Yet granulocyte proportions did not demonstrate any concordance between methods (ρ = 0, *p* = 1). Lymphocyte proportions were more likely to be larger from flow cytometry (67%), whereas most granulocyte proportions were overestimated through traditional hematology compared to flow cytometry (93%).

**Figure 3.**
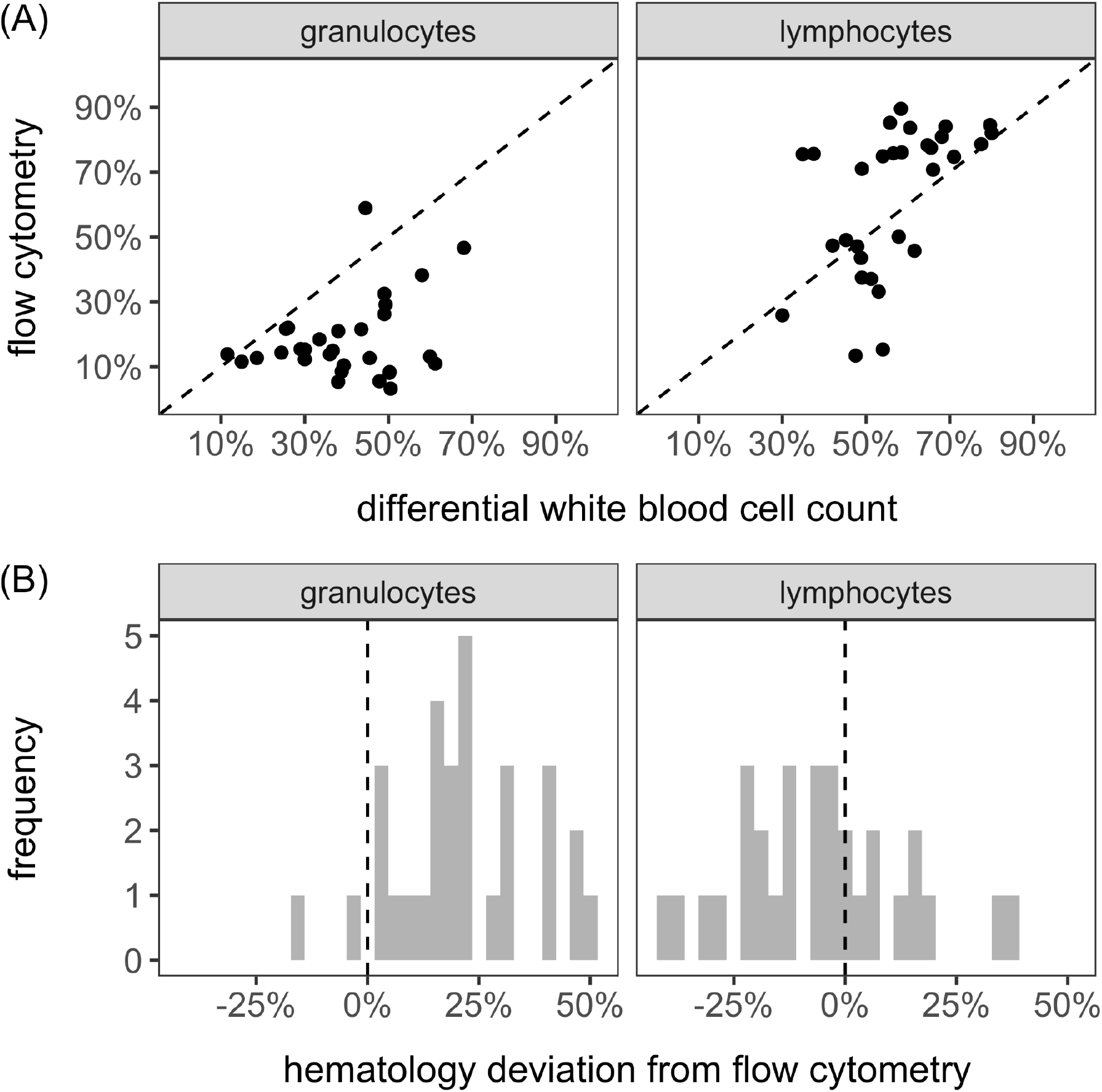
(A) Individual-level relationship between manual differential white blood cell counts and cells identified through flow cytometry for granulocyte and lymphocyte proportions. The dashed line shows the 1:1 correspondence between methods. (B) Deviations from the 1:1 line, where larger values indicate hematology overestimating cell proportions from flow cytometry.

GAMs revealed seasonal trends in most immune cell populations identified by flow cytometry, after adjusting for precision covariates (Table 2). Granulocyte populations marginally decreased across the Oklahoma occupancy period, whereas lymphocyte populations significantly increased from spring migration through the reproductive season and fall migration (Figure 4).

**Table 2.**
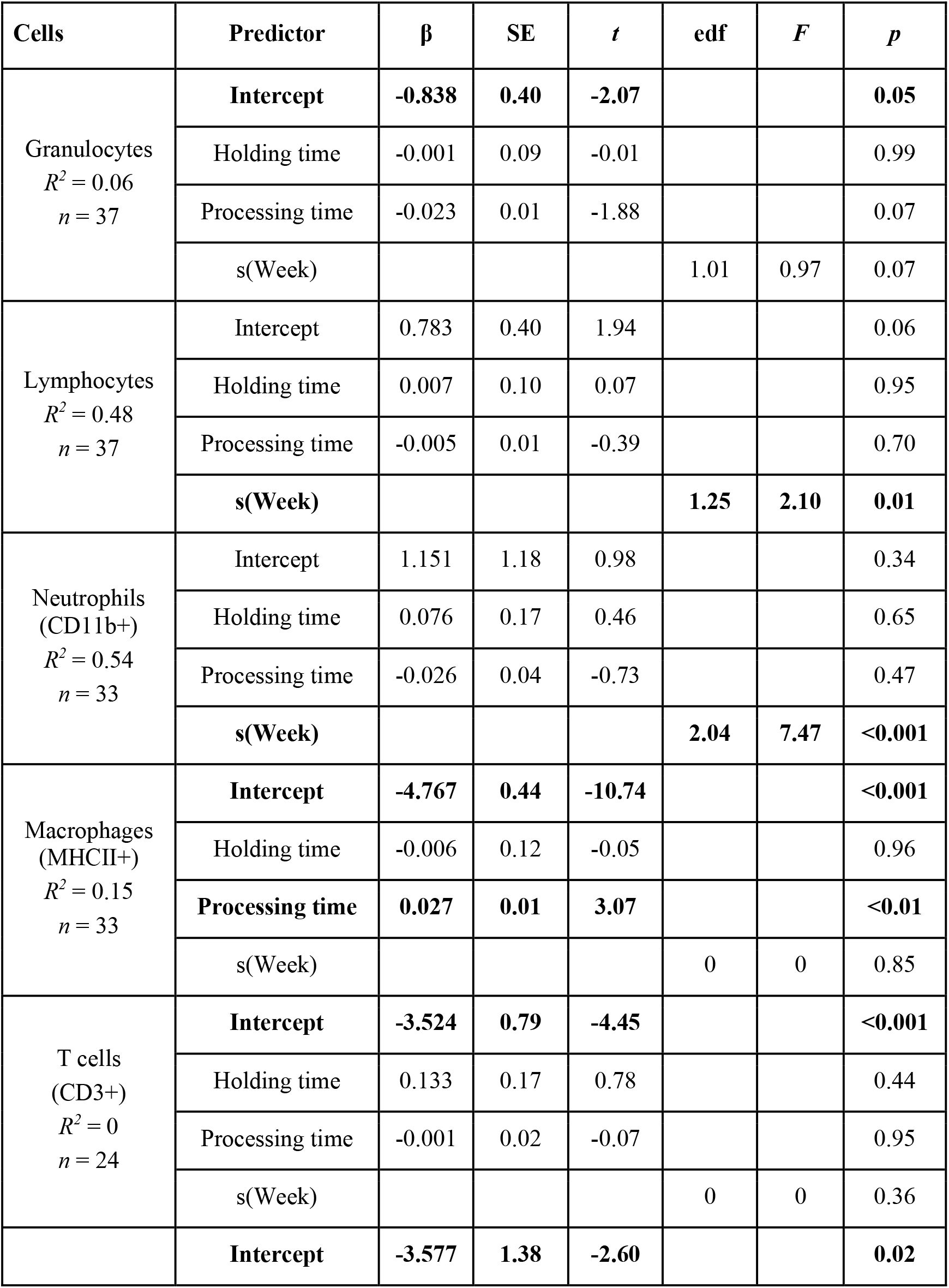
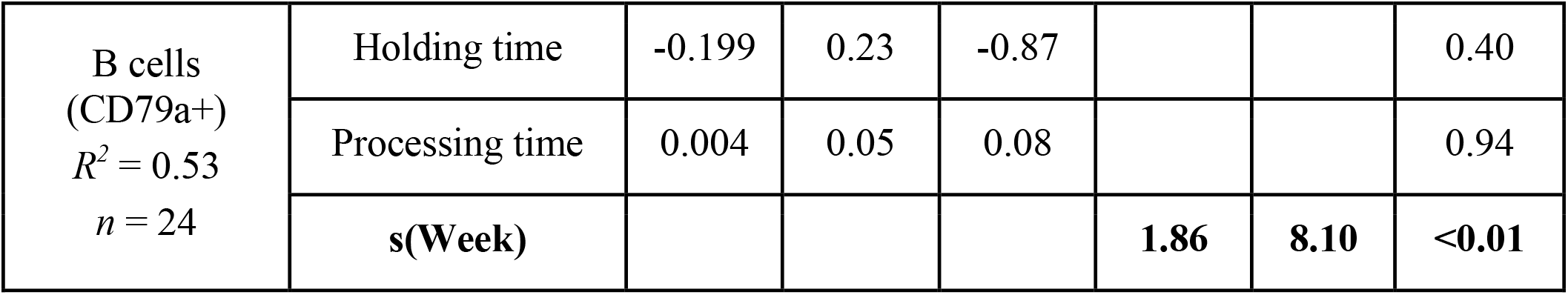
Estimated fixed effects and smoothed terms from immune cell proportion GAMs (quasibinomial response). Statistically significant effects are displayed in bold font.

**Figure 4.**
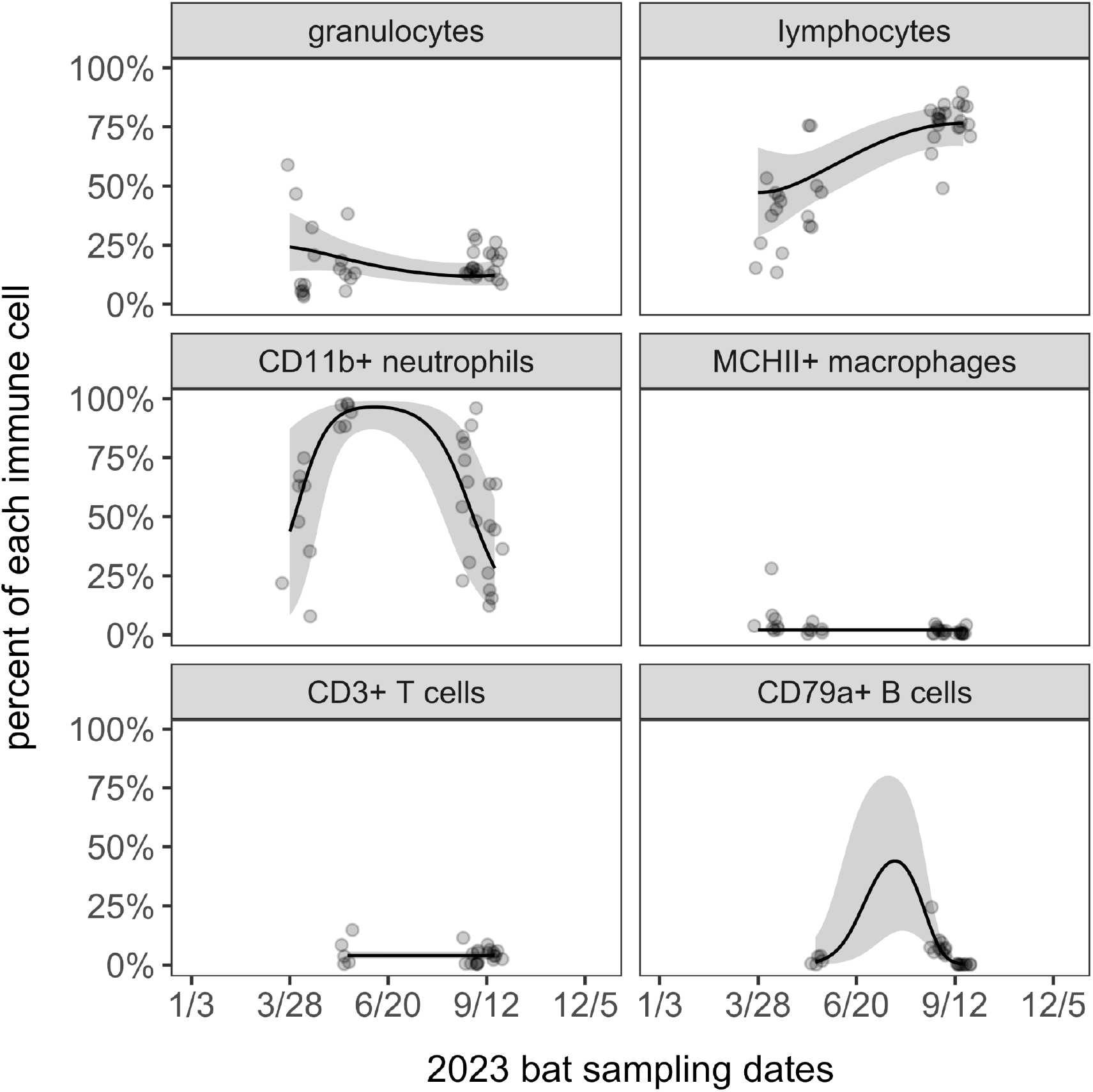
Seasonal patterns in the innate (granulocytes) and adaptive (lymphocytes) branches of the Mexican free-tailed bat cellular immune system. Raw data are jittered to reduce overlap and overlaid with respective GAM fits (quasibinomial response) and 95% confidence intervals.

For specific immune cell subsets (Figure 4), CD11b+ neutrophils were most abundant during the reproductive period and were lowest during migratory periods, whereas CD79a+ B cells were most abundant following reproduction and as pups became independent. By contrast, neither MHCII+ macrophages nor CD3+ T cells showed any seasonal variation. Analogous GAMs that tested for sex-biased cell populations found no significant differences, with the exception of females having greater proportions of CD79a+ B cells than males (Table S1).

We detected no significant trade-offs between or within different branches of the cellular immune system, although effect sizes suggested weak but consistently negative associations for granulocytes and lymphocytes (ρ = -0.08, *p* = 0.63), CD11b+ neutrophils and MHCII+ macrophages (ρ = -0.17, *p* = 0.35), and CD3+ T cells and CD79a+ B cells (ρ = -0.22, *p* = 0.29).

## Discussion

Characterizing how seasonal patterns in wildlife immunity relate to energetic trade-offs can improve our ability to predict when hosts are likely to shed pathogens, informing strategies to prevent or manage infectious disease risks (Martin et al. 2008; Plowright et al. 2024). Yet for priority taxa such as bats, our ability to profile the immune system in wild hosts, and especially cellular defenses, has been limited (Schountz 2014; Banerjee et al. 2018). Here, we adapted flow cytometry protocols to wild bat systems and tested seasonal fluctuations in specific immune cells in the context of migration and pregnancy. We validated the use of small blood volumes typically obtained in field studies of many bats and showed that antibodies previously used to identify immune cells in Egyptian fruit bats are largely cross-reactive in the Mexican free-tailed bat. Flow cytometry also outperformed traditional hematology in quantifying bat leukocyte profiles. In turn, we found distinct seasonal changes in immune cell populations of Mexican free-tailed bats between spring and fall migration, providing insight into how phenology influences cellular immunity. These findings provide proof of concept for applying flow cytometry to field studies of wildlife and provide novel insights into the seasonality of wild bat cellular immunity.

While flow cytometry can serve as a powerful tool for characterizing wildlife cellular immunity, the limited availability of reagents for non-model organisms requires first identifying cross-reactive antibodies (Moreira et al. 2015; Migalska et al. 2023). However, determining cross-reactivity can be challenging, as antigenic similarity does not always equate to functional binding for a given antibody. As such, this process typically relies on a combination of sequence homology, positive controls, and testing across target taxa to confirm antibody performance, although single-cell RNA sequencing is increasingly detecting conserved genes that can enable later generation of monoclonal antibodies (Li et al. 2022; Chen et al. 2024; Watson et al. 2024). Previously, there were no antibodies confirmed for Mexican free-tailed bats, and those tested here were earlier validated using initial single-cell RNA sequencing in Egyptian fruit bats (Friedrichs et al. 2022). The genera *Rousettus* and *Tadarida* represent deeply diverged bat lineages that last shared a common ancestor 50–60 million years ago (Teeling et al. 2005). This divergence makes antibody cross-reactivity uncertain, as epitope sequences and protein structures can vary widely across such evolutionary distances (Brodsky and Parham 1982; Brodersen et al. 1998; Conrad et al. 2007). Interestingly, four of the five monoclonal antibodies tested were compatible with Mexican free-tailed bats, suggesting some of these antibodies may be based in highly conserved mechanisms that span the Chiroptera. This apparent conservatism could allow broader applicability of these antibodies to be cross-reactive in other bat lineages (Becker et al. 2025c). However, these results should be interpreted as preliminary, as additional validation across diverse bat species will be necessary to confirm epitope homology and assess how consistently antibody binding would occur beyond the single species examined in this study.

Flow cytometry provided a more specific and, in some cases, more accurate approach to quantifying immune cell composition compared to differential white blood cell counts. When comparing broad cell groups identified by both approaches, we found greater concordance between methods for lymphocyte counts, whereas granulocyte counts showed greater disparities. The direction of both discrepancies suggested manual hematology likely underestimated the relative abundance of lymphocytes while overestimating the relative abundance of granulocytes. Such differences may be driven in part by our identification of only 100 leukocytes, which is a common approach to differential counts and balances quantification with the time-intensive nature of manual analysis (Siekmeier et al. 2001; Bain et al. 2006; Davis et al. 2008). Yet this limitation can lead to rare cells such as eosinophils and basophils being underrepresented, a bias that is only partly corrected by manually identifying more leukocytes (Weisbrod et al. 2021).Improved ability to quantify rarer immune cells, and to distinguish between subsets of broad cell categories (e.g., not only lymphocytes but T and B cells specifically) is especially important in the context of intra- and interspecific comparisons. By enabling consistent cell identification and quantification using conserved markers, flow cytometry offers a much-needed tool for improving the comparative study of immunity in wild hosts and standardizing data across diverse systems.

The application of flow cytometry to our migratory bat system primarily supported our hypothesis of favoring investment toward innate immune cells during energetically costly periods of migration and reproduction, given the lower developmental costs of these defenses (Rauw 2012; McDade et al. 2016). Lymphocyte counts were lowest following spring migration and during pregnancy, increasing during the non-reproductive period and prior to fall migration; we observed inverse trends, albeit weaker in strength, for granulocyte counts. Such patterns could indicate greater energetic costs of having undertaken spring migration than of preparing for fall migration, particularly given the coupling of migratory arrival into Oklahoma with early pregnancy (Bernardo and Cockrum 1962; McGuire and Guglielmo 2009). Such findings also agree with previous work using differential white blood cell counts in other migratory bat species, in which Nathusius’ pipistrelles (*Pipistrellus nathusii*) had more lymphocytes and fewer neutrophils in fall migration compared to pre-migratory periods and in which some migratory strategies in silver-haired bats (*Lasionycteris noctivagans*) were associated with neutrophilia after spring migration (Voigt et al. 2020; Rogers et al. 2022). However, sampling logistics limited our ability to fully profile cellular immunity throughout the reproductive season (e.g., lactation) and during fall migration itself (i.e., late September, early October). As such, more fine-scale seasonal sampling, with annual replication, would more fully characterize tradeoffs between bat energetics and different branches of cellular immunity. Despite this limitation, our validation of additional antibodies allowed us to also query seasonal shifts within adaptive immunity, finding elevated B cells after reproduction and before fall migration but no change in T cells. While this pattern was restricted by small sample size, shifts in B cells could be relevant for understanding the seasonality in bat viral infections (Lam et al. 2020; Roffler et al. 2024).

Flow cytometry has been applied to a small subset of bats (Martínez Gómez et al. 2016; Periasamy et al. 2019; Gamage et al. 2020; Friedrichs et al. 2022; Chen et al. 2024), a diverse order of over 1,500 species with substantial immunological variation (Becker et al. 2025c). Here, we expand the application of flow cytometry into a distinct bat lineage (i.e., Molossidae) and, critically, into the context of wild hosts. Although our work provides proof of concept and new insights into immune cell seasonality of Mexican free-tailed bats, we encourage greater adoption of flow cytometry into other wild systems, where factors such as land conversion, urbanization, and hibernation may shape bat cellular immunity (Whiting-Fawcett et al. 2021; Tovstukha et al. 2025; Becker et al. 2025a). Similarly, overcoming the challenges posed by adopting flow cytometry to studies of wildlife could benefit other small vertebrate systems. In particular, studies of wild rodents could capitalize on monoclonal antibodies generated for laboratory mice and rats, given the conserved nature of many cell markers (Jones et al. 1993; Kühnlein et al. 1996; Migalska et al. 2023). More broadly, accurately quantifying immune cell populations in wild hosts in the context of intrinsic and extrinsic stressors could help predict zoonotic risks, minimize human–wildlife conflict, and guide conservation strategies (Ohmer et al. 2021; Simonis et al. 2025). By advancing these specific techniques, we can continue to refine our approach to wildlife immunology, ultimately aiding in conservation and public health efforts.

## Ethical approval

Field procedures were performed according to guidelines for the safe and humane handling of bats from the American Society of Mammalogists (Sikes and Gannon 2011), approved by the Institutional Animal Care and Use Committee of OU (2022-0198), and permitted by the Oklahoma Department of Wildlife Conservation (10567389) and Oklahoma State Parks.

## Acknowledgements

We thank the Oklahoma Department of Wildlife Conservation and Oklahoma State Parks for cave access and the Selman Living Laboratory and University of Central Oklahoma for logistical support. We also thank Jim Henthorn and the Institutional Research Core Facility at the University of Oklahoma Health Campus for the use of the Flow Cytometry and Imaging facility. We also thank Molly Simonis for fieldwork support and Hayley Lanier for manuscript feedback. Lastly, we thank the late William Caire for introducing us to the western Oklahoma cave sites.

This work was supported by the National Institute of General Medical Sciences of the National Institutes of Health (P20GM134973) and Research Corporation for Science Advancement (RCSA; subaward 29018, part of a USDA Non-Assistance Cooperative Agreement with RCSA Federal Award 58-3022-0-005). Support was also provided by the Oak Ridge Associated Universities (Ralph E. Powe Junior Faculty Enhancement Award) and the University of Oklahoma (Vice President for Research and Partnerships and Data Institute for Societal Challenges). DJB was also supported by the Edward Mallinckrodt, Jr. Foundation, National Science Foundation (DBI 2515340), and National Institutes of Health (R21AI190246).

## Notes

### Competing Interest Statement

The authors have declared no competing interest.

