## Supplemental Material for "Adapting flow cytometry for studying immune cell seasonality in wild migratory bats"

Figure S1. Immune cell proportions as a function of blood volume from laboratory mice

Table S1. Immune cell proportions in Mexican free-tailed bats as a function of sex

Figure S1. Populations of CD4 T cells, CD19 B cells, and neutrophils as a function of blood sample volume in laboratory mice ($n = 2$). Plots show means with corresponding standard error.

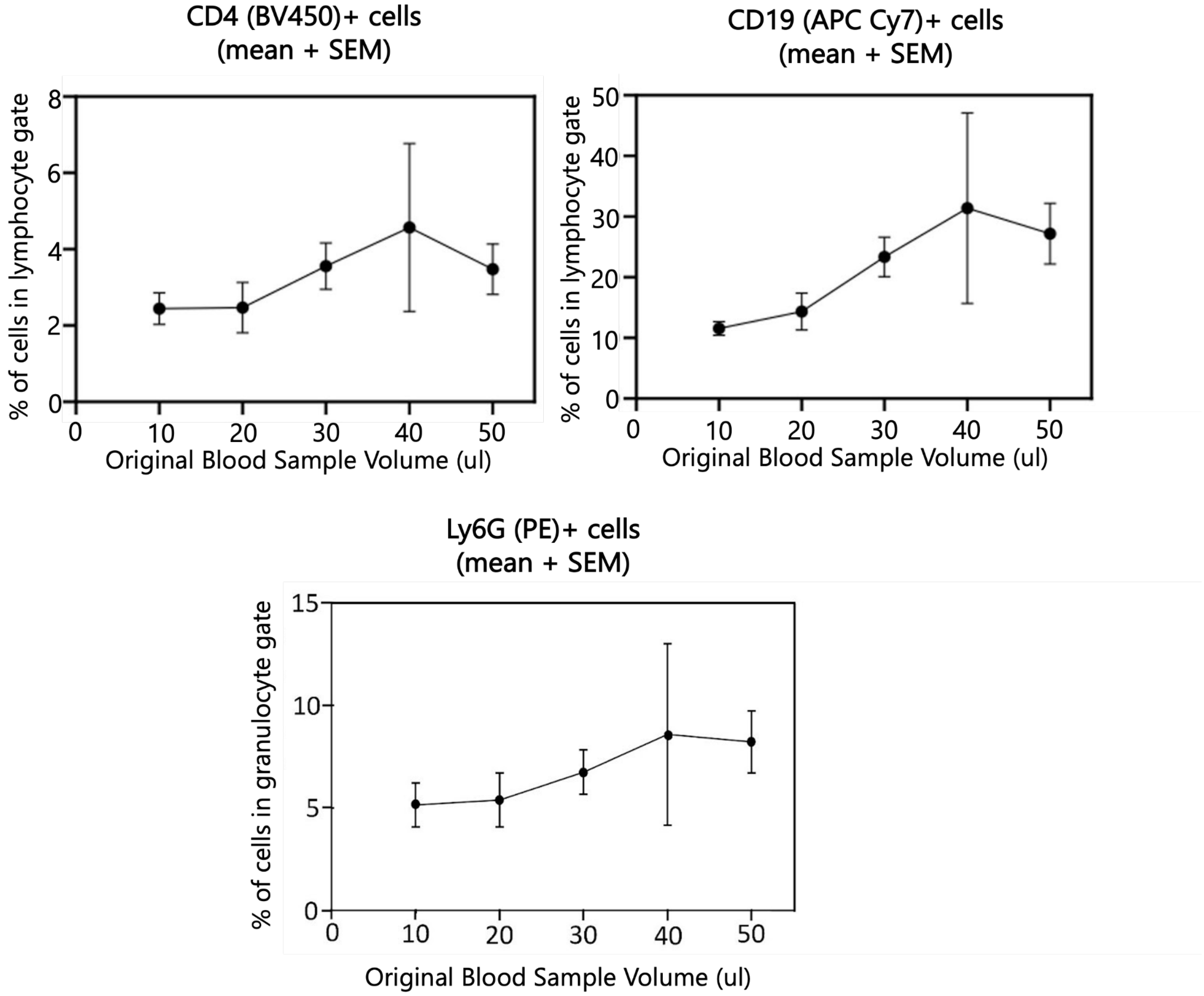

Table S1. Estimated effects from immune cell proportion models (quasibinomial response) with sex as the main predictor, accounting for holding time and processing time ($p \leq 0.05$ in bold).

| Cells | Predictor | β | SE | *t* | *p* |
|---|---|---|---|---|---|
| Granulocytes $R^2 = 0$ | **Intercept** | **-1.374** | **0.40** | **-3.45** | **<0.01** |
| | Male | 0.058 | 0.21 | 0.27 | 0.79 |
| | Holding time | -0.032 | 0.09 | -0.35 | 0.73 |
| | Processing time | -0.007 | 0.01 | -0.95 | 0.35 |
| Lymphocytes $R^2 = 0.35$ | **Intercept** | **1.527** | **0.43** | **3.54** | **<0.01** |
| | Male | -0.109 | 0.24 | -0.46 | 0.65 |
| | Holding time | 0.098 | 0.10 | 0.99 | 0.33 |
| | **Processing time** | **-0.031** | **0.01** | **-4.75** | **<0.001** |
| Neutrophils (CD11b+) $R^2 = 0.14$ | Intercept | 0.993 | 0.92 | 1.08 | 0.29 |
| | Male | -0.169 | 0.45 | -0.37 | 0.71 |
| | **Holding time** | **-0.403** | **0.20** | **-2.05** | **0.05** |
| | Processing time | 0.025 | 0.02 | 1.34 | 0.19 |
| Macrophages (MHCII+) $R^2 = 0.12$ | **Intercept** | **-4.663** | **0.64** | **-7.24** | **<0.001** |
| | Male | -0.080 | 0.35 | -0.23 | 0.82 |
| | Holding time | -0.023 | 0.15 | -0.16 | 0.87 |
| | **Processing time** | **0.026** | **0.01** | **2.92** | **<0.01** |
| T cells | **Intercept** | **-3.693** | **0.98** | **-3.76** | **<0.01** |

| | | | | | |
|---|---|---|---|---|---|
| (CD3+) $R^2 = 0$ | Male | 0.132 | 0.43 | 0.31 | 0.76 |
| | Holding time | 0.157 | 0.19 | 0.83 | 0.42 |
| | Processing time | 0.00 | 0.02 | -0.01 | 0.99 |
| B cells (CD79a+) $R^2 = 0.47$ | **Intercept** | **3.075** | **1.37** | **2.24** | **0.04** |
| | **Male** | **-1.260** | **0.51** | **-2.48** | **0.02** |
| | **Holding time** | **-1.122** | **0.29** | **-3.89** | **<0.001** |
| | **Processing time** | **-0.138** | **0.05** | **-3.06** | **<0.01** |
